# The Innexin 7 gap junction protein contributes to synchronized activity in the *Drosophila* antennal lobe and regulates olfactory function

**DOI:** 10.1101/2025.01.19.633780

**Authors:** Nicolás Fuenzalida-Uribe, Sergio Hidalgo, Bryon Silva, Saurin Gandhi, David Vo, Parham Zamani, Todd C. Holmes, Sercan Sayin, Ilona Grunwald-Kadow, Dafni Hadjieconomou, Diane K. O’Dowd, Jorge M. Campusano

## Abstract

In the mammalian olfactory bulb (OB), gap junctions coordinate synchronous activity among mitral and tufted cells to process olfactory information. In insects, gap junctions are also present in the Antennal Lobe (AL), a structure homologous to the mammalian OB. The invertebrate gap junction protein ShakB contributes to electrical synapses between AL Projections Neurons (PNs) in *Drosophila*. Other gap junction proteins, including Innexin 7 (Inx7), are also expressed in the *Drosophila* AL, but little is known about their contribution to intercellular communication during olfactory information processing. Here we report spontaneous calcium transients in PNs grown in cell culture that are highly synchronous when these neurons are physically connected. RNAi-mediated knock down of Inx7 in cultured PNs blocks calcium transient neuronal synchronization. *In vivo*, downregulation of Inx7 in the AL impairs both vinegar-induced electrophysiological calcium responses and behavioral responses to this appetitive stimulus. These results demonstrate that Inx7-encoded gap junctions functionally coordinate PN activity and modulate olfactory information processing in the adult *Drosophila* AL.

## INTRODUCTION

The organization and operation of olfactory circuits are highly conserved between vertebrates and invertebrates. Olfactory information is first received by the Olfactory Receptor Neurons (ORNs) that project to discrete glomerular structures in the Olfactory Bulb (OB) in mammals and the Antennal Lobe (AL) in insects. Within glomeruli, olfactory information is integrated and then transmitted to higher processing centers, including cortical areas in mammals and the Mushroom Bodies (MBs) in insects, where olfactory memories can be formed and stored [1–4]. In addition to the anatomical organization, the similarities between vertebrate and invertebrate olfactory systems extend to the cellular and molecular mechanisms responsible for processing odorant information. These cellular and molecular olfactory mechanisms, including those used in invertebrate models, are not completely understood,.

Electrical synapses, formed by gap junctions, are important players in neuronal communication, information processing, and ultimately brain function [5, 6]. This type of cell communication underlies most of the coupled activity observed within neural circuits, including the mammalian olfactory system. For instance, the synchronized activity recorded between the mitral and the tufted cells of the OB glomeruli, which is involved in the processing of olfactory information and stimuli gain, is mediated by electrical synapses [7–10]. Likewise, electrical synchronization within the AL is also thought to be important in pheromone and food-related odor processing in insects [11–14].

Gap junctions are formed by specific proteins: in vertebrates, they are encoded by two gene families, connexins and pannexins, while in invertebrates, gap junctions are formed by just one family of proteins, the innexins [15, 16]. Out of the eight innexins identified in the *Drosophila melanogaster* genome (*inx 1-8*), four of them (*inx5*, *inx6*, *inx7*, and *shakB*, also known as *inx8*) show detectable mRNA levels in neurons in the pupal and adult fly brain. Notably, *lnx7* and *shakB* innexins exhibit the highest transcript levels [17]. Consistent with these expression studies, it has been shown that *shakB* plays a role in developing *Drosophila* optic lobe lamina [18]. Importantly, ShakB shows the highest expression in mature fly AL [17, 19], which is consistent with the finding that this innexin mediates electrical synapses between sister PNs and also between pairs of AL lateral neurons (LNs) [20–22]. Less is known regarding the contribution of other innexins to brain development or function. In this regard, it was demonstrated that Inx7 plays a role in axon guidance and nervous system development [23]. In addition to this, it was reported that Inx7 forms heterotypic gap junctions between two modulatory neurons in MB that contribute to olfactory memory [24].

Here we focus on the functional role that Inx7 plays in *Drosophila* olfactory function. Our data demonstrate that knock-down of Inx7 in AL PNs decouples correlated calcium activity between physically connected PNs grown in cell culture. *In vivo*, evoked calcium signals in *Drosophila* AL by vinegar exposure are impaired by knock down of Inx7 in this fly brain region. Furthermore, we show that Inx7 expression in AL PNs is required for olfactory behavioral responses to vinegar exposure. These results reveal a previously unrecognized role of Inx7 gap junctions to the integration of olfactory sensory information in flies

## MATERIALS AND METHODS

### Flies

To identify antennal lobe PNs, flies bearing a Gal4 transgene expressed in AL PNs (*w^1118^*;*GH146-Gal4* [25]), were crossed to flies containing a UAS-GFP transgene. Strains were obtained from the Bloomington *Drosophila* Stock Center (BDSC, Indiana University, IN) (strains # 30026 and 1521, respectively). In F1 of this cross it is possible to detect GFP expression in the AL (see Figure 1A). Recombinant animals homozygous for these two transgenes were obtained (from here on GH146,GFP), and were crossed to a strain that allows the expression of an RNAi targeting Inx7 (RNAi*^inx7^*) under the control of UAS (*w^1118^*;*UAS-RNAI^inx7^*), which was obtained from the Vienna *Drosophila* Resource Center, VDRC, Vienna, Austria) (strain ID#22949, that we called Inx7-KD-a-PN). A second RNAi*^inx7^*strain was also used for some preliminary experiments (VDRC #22948). In some experiments (see below), *elav-gal4*,*w^1118^* animals were used to drive the RNAi*^inx7^*. Flies were kept at 18°C throughout their development to diminish the expression of gal4-driven genes. Only 3 days before they were used for experiments they were brought to room temperature (22°C). For additional experiments we used the strain ID#50856, obtained from BDSC.

**Figure 1.**
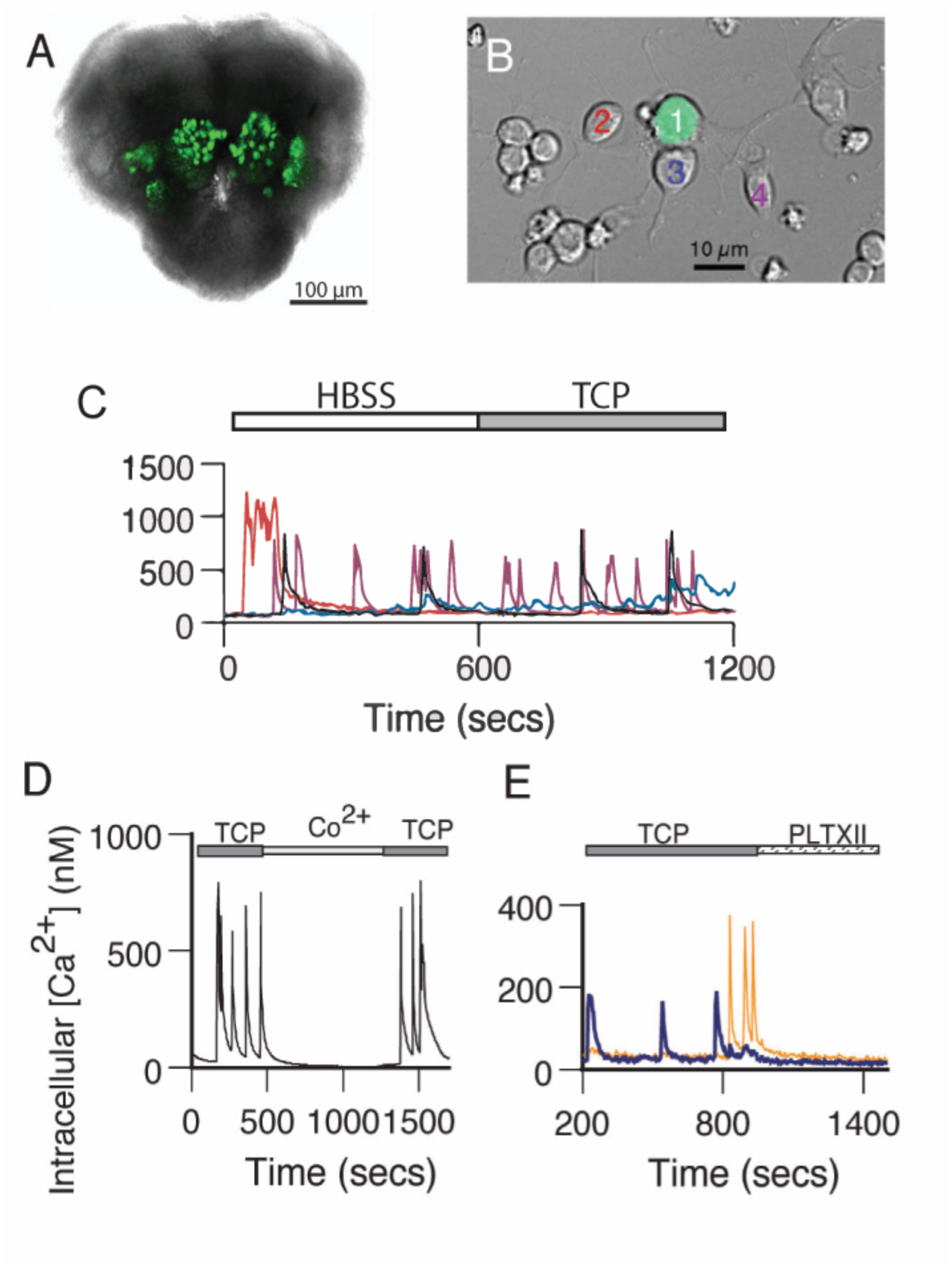
GFP-positive PNs generate spontaneous calcium transients independent of fast chemical synaptic transmission. **A.** GFP-positive antennal lobe PNs in the brain of a late-stage pupa. Bright-field image with a fluorescent overlay of the anterior face in a brain from a GH146>GFP pupa at ∼72 hrs after pupation. **B.** GFP-positive PN surrounded by GFP-negative neurons in dissociated cell culture at 3 DIV. Fluorescent mask projected on a Nomarski image. **C.** Spontaneous calcium transients in GFP-positive PNs recorded in standard saline (HBSS) are not affected by addition of TCP, a cocktail of drugs that block synaptic transmission (1µM TTX, 20 µM curare, 10 µM picrotoxin). Traces correspond to the calcium levels in the neurons presented with different colors in B. **D.** Addition of cobalt (Co^2+^, 2 mM) reversibly blocks spontaneous calcium transients. Representative experiment. **E.** Exposure to an irreversible voltage-gated calcium channel blocker, Plectreurys toxin (PLTXII, 50 nM) inhibits spontaneous calcium transients. In this and all other subsequent figures, intracellular calcium levels were monitored using the ratiometric imaging dye Fura-2 AM. The 340/380 ratios were sampled at 4-second intervals and converted to estimates of intracellular calcium based on a standard calcium curve.

### Quantitative PCR experiments

The efficacy of the RNAi*^inx7^* line was assessed by qPCR. For these experiments, RNA was prepared from adult fly heads obtained after elav-Gal4,w*^1118^* flies were mated with *w^1118^*;UAS-RNAi*^inx7^* animals. RNA obtained from the heads of parental strains was used as control.

The procedure to carry out PCR was as described [26]. Briefly, cDNA was prepared from each RNA sample with 2 μg of RNA as starting material. RT-qPCR was carried using the 5 × HOTFIREPol® EvaGreen® qPCR Mix Plus Kit (Solis BioDyne) and LightCycler® (Roche) apparatus. PCR parameters were: 94°C for 6 minutes, followed by 40 cycles of 94 °C for 10 s, 56°C for 15 s and 72°C for 20 s. Primer efficiency was evaluated after a serial dilution of the reaction mix and was calculated as 10^−1/slope^. The *ribosomal protein 49* gene was used for RNA quantitation. Primer pairs were designed with MacVector software: RP49F1, 5’-caagggtatcgacaacagagtcg-3’; RP49B1, 5’- tgcaccaggaacttcttgaatcc-3’ (amplicon of 126 bp), and *inx7*F9, 5’-tttggtgactctggctccattc– 3’, 5’-cgcaggaagaaaaggaacatcc–3’ (amplicon of 287 bp). Analysis of relative gene expression (*inx7* target gene versus *RP49* reference gene) in knockdowns vs control lines was calculated according to the 2^-ΔΔCT^ method.

Our results show that the two RNAi strains showed differential efficiency at decreasing *inx7* transcript expression: the ID#22949 resulted in a decrease of 85 ± 7% of *inx7* mRNA expression, while the ID#22948 second RNAi strain reduced *inx7* transcript expression to only 50 ± 7% (n= 3 independent experiments per strain). Thus, we decided to work only with the RNAi*^inx7^*ID#22949.

### Primary neuronal cultures

Primary neuronal cultures were prepared as previously described [26, 27]. In initial studies, dissociated neuronal cultures were prepared with one pupal brain per culture. Given the small number of PNs found in one brain, cultures contained only a small number of GFP-positive PNs and a much larger population of GFP-negative neurons. The number of GFP-positive PNs counted in 3-4 days *in vitro* (DIV) cultures was 11.9 ± 1.6 per brain (mean ± SEM, n=39 cultures). This represents a mean recovery rate (assuming ∼100 GFP-labeled PNs/brain) of about 10-15%. To increase the number of GFP-expressing neurons in our preparation, all cultures were prepared from the equivalent of 2 brains/culture. Cultures were maintained in a humidified 5% CO2 incubator at 22-24°C and imaging experiments were performed on cultures 4 to 8 DIV.

### *In vitro* calcium imaging

Calcium imaging studies were performed as previously reported [27, 28]. Briefly, cultures were loaded with the calcium indicator dye Fura2-AM (5 mM, Molecular Probes, Eugene OR), in presence of pluronic acid (0.1%, Invitrogen), in a HEPES-buffered salt solution (HBSS). Cultures were imaged on an inverted Nikon microscope with a 40x, 1.3 numerical aperture, oil immersion objective. Fluorescent illumination was provided by a 150-W Xenon arc lamp and filters at 340 and 380 nm were used for excitation. All images were acquired at the emission wavelength 510 nM and recorded every 4 sec via a digital CCD Camera (Photometrics, Tucson, AZ). Pseudocolor 340/380 ratio images were generated with Metafluor software version 5.1 (MDS Analytical Technologies, Sunnyvale, CA). Ratiometric values were converted to estimates of intracellular calcium concentration.

Prior to beginning an experiment each culture was scanned to determine if they contained at least two GFP-positive PNs in close proximity (pairs). PN pairs, either with adjacent cell bodies or overlapping neuritic processes were found in ∼1 out of 5 cultures and these were selected for calcium imaging. Intracellular calcium levels were evaluated by determining the average signal over neuronal somata. Neurons were included in the data set if a stable recording was maintained for at least 10 minutes, and an increase in the 340/380 ratio greater than 600 nM was induced by exposure to 5 µM ionomycin at the end of the experiment. A calcium transient was defined as an increase in intracellular calcium concentration greater than 60 nM (>5 times baseline noise level) that reaches its peak within 20 seconds and declines by at least 50% from the peak within 3 minutes. The transient frequency in each cell was calculated as the number of transients divided by the time of recording. The amplitude of each transient was measured from baseline to peak and an average value for each cell was calculated [29]. When mentioned, TCP solution was used to block fast chemical neurotransmission. TCP consisted of <u>T</u>TX (1 µM, Alomone Labs, Israel, to block voltage-gated sodium channels); <u>C</u>urare (d-tubocurarine, 20 µM, Sigma, to block nAChRs); and <u>P</u>icrotoxin (PTX, 10 µM, Sigma, to block GABAA receptors). The following drugs were bath applied in specific imaging experiments: Plectreurys toxin (PLTX-II, 50 nM, Alomone Labs, Israel), cobalt chloride (Co^2+^, 2 mM, Sigma).

A cross-correlation function (CCF) was used to evaluate the correlation in time of spontaneous calcium transients in PN pairs in which both cells were active (>80% of pairs examined). The CCF was evaluated for a time-window of at least 10 minutes, using the routine available in Clampfit9 (Molecular Devices, CA): the data set of one cell was shifted along the second, one sample at a time, in both directions. Thus, the CCF measured the correlation between data sets at each time point. The correlation coefficients reported here are the value of the CCF at lag time 0.

### *In vivo* calcium imaging

Female flies (4- to 7-day-old) of the genotype GH146-Gal4,UAS-GCaMP3 were used for these experiments. In vivo imaging was made as previously [30]. All imaging experiments were conducted using a Leica DM6000FS fluorescent microscope, a 40x water immersion objective, and a Leica DFC360 FX fluorescent camera, and focusing on the AL plane. Images were acquired using the Leica LAS AF E6000 software for a total recording time of 60s, at a rate of 30 frames per second with 4 × 4 binning mode. Time series images were acquired at 256 × 256 pixel resolution at 30 frames per second. For odor delivery, we used a Syntech stimulus controller CS-55 (Syntech) and mass flow controllers. Throughout the experiments, a charcoal-filtered continuous humidified airstream (1L/min) was delivered through an 8-mm Teflon tube positioned ∼10 mm away from the fly antenna. For odor stimulation, odors were delivered into the main airstream by redirecting ∼30% of the main airflow for 1 s through a head-space glass vial containing appropriately diluted odorant. Balsamic vinegar (Alnatura) was diluted in distilled water. To measure the fluorescent intensity change, a region of interest (AL glomeruli) was drawn manually, and the resulting time trace value was used for data analysis. The relative change in fluorescence intensity was calculated by using the following formula: ΔF/F = 100(F_n_ − F_0_)/F_0_, where F_n_ is the n^th^ frame after stimulation, and F_0_ is the average basal fluorescence of 5 frames before stimulation. All data were normalized to background air. For data analysis, we used the peak maximum value of the response to stimulation. The pseudocolored images were generated using ImageJ. All data processing and statistical tests were done using Excel and GraphPad Prism softwares, respectively.

### Spherical Treadmill behavioral assay

The spherical treadmill assay, the online speed data acquisition rate, and offline calculations were performed as reported [31]. In brief, a tethered female fly was transferred onto the treadmill and assayed after a 3-min period of acclimatization. The assay consisted of 10 consecutive trials of vinegar odor exposure, which were separated by intervals of 60 s. A custom-made PTFE (Teflon) with a 4mm diameter tube, fixed at ∼3mm distance from the tethered fly, was used for the delivery of the appetitive olfactory stimulus. The air speed was set to 100 mL/min via a Natec Sensors mass-flow controller. The balsamic vinegar solution was prepared daily at 20% v/v dilution in 100 ml Schott bottles. Each trial was recorded for a minimum of 52 s. The recording was divided into pre-stimulation (20 s), stimulation (12 s), and post- stimulation (30 s) periods and the recorded speed data were downsampled to 10Hz by summation. The average run activity was measured as the fraction of time where flies showed running speed higher than 0mm/s while being stimulated with the odor. A stop was defined as flies not moving (0mm/s) for at least 100ms. Data was visualized with matplotlib (1.4.2) and graphed as average run speed (mm/s).

### Olfactory preference test

Flies were placed in starvation conditions (vial without food, with a filter paper band moistened with water) at 25°C for 24 hours before the beginning of an experiment. Groups of 30-50 flies (four to six days old), were assessed in the olfactory acuity assay, which was carried out as previously described [26]. The behavior room was kept at 23-26°C with 60-70% relative humidity under dim red light. We used a response index (RI) to express a choice preference for the vinegar odor, calculated by subtracting the number of flies in the air arm from the number of flies in the vinegar arm, divided by the total number of flies. RI equals 1 would indicate 100% preference for the vinegar odor.

### Single-locomotor analysis

As previously reported [32, 33]. Briefly, single flies were placed on a circular white arena (39 mm diameter, 2 mm high) to be video recorded for 3 minutes at room temperature. Videos were analyzed offline using a video tracking analysis (Buridan-tracker).

## RESULTS

### Pairs of PNs show correlated calcium transients that depend on Inx7 expression

Spontaneous calcium activity has been previously reported in cultured *Drosophila* neurons [28, 29]. To examine whether cultured AL PNs also show spontaneous activity, we monitored calcium concentration under basal conditions using the ratiometric intracellular calcium indicator, Fura-2. AL PNs, identified by cell-specific GFP expression, show calcium transients (Figure 1A-C) like those observed in Kenyon cells (KCs) [28, 29]. Calcium signals vary in frequency and waveforms and are not exclusive to AL PNs, as surrounding (GFP-negative) neurons also display these transients (Figure 1C).

To characterize the nature of the spontaneous calcium transients observed in AL PNs in culture, we performed recordings under different conditions (Figure 1C-E). The calcium transients observed in PN are still present upon exposure to fast-synaptic blockers (TTX, Curare, and Picrotoxin; TCP), suggesting that they are independent of chemical synapses (Figure 1C). Bath application of either Co^2+^ or PLTXII, an insect-specific calcium channel antagonist [28, 29], completely abolishes these spontaneous events. These data indicate that calcium transients depend on the activity of PLTXII- sensitive voltage-gated calcium channels (Figure 1D-E).

Although the AL PNs represent a small subset of the number of the total cell population in culture, from time to time we observed pairs of GFP-positive neurons in direct physical contact (Figure 2A-C). The calcium transients in most of these pairs show a high degree of correlation over time under basal recording conditions (Figure 2A-A’).

**Figure 2.**
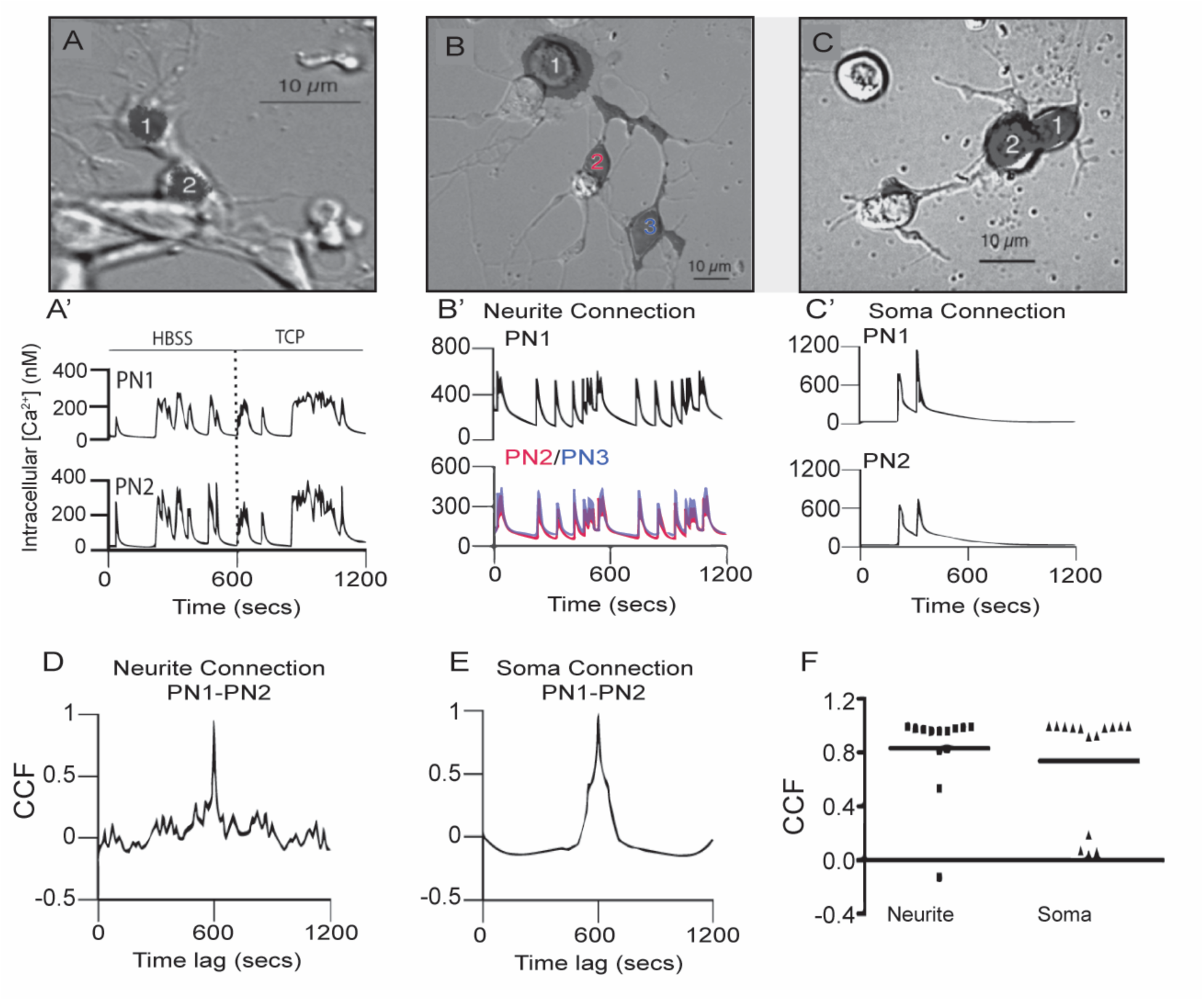
Spontaneous calcium transients are highly correlated in time in PN-PN pairs connected by neurites or cell bodies. **A.** A pair of physically connected GFP- positive PNs in dissociated cell culture at 5 DIV. Fluorescent mask (shaded black) projected on Nomarski image. **A’**. PN 1 and 2 generate spontaneous calcium transients that are highly correlated in time in HBSS. Correlated calcium transients in the pair persist following the addition of TCP to block fast chemical synaptic transmission. **B.** Three PNs in contact through neuritic processes that were regenerated in culture. **B’.** PN1, 2 and 3, connected by newly regenerated neurites (PN2 in red and PN3 in blue, panel B), have spontaneous calcium transients that are highly correlated in time. **C.** Two PNs’ in contact at their cell bodies. Fluorescent masks (shaded black) projected on Nomarski images. **C’.** Correlated spontaneous calcium transients in PN1 and PN2 (panel C) in contact at their somas. CCF for traces shown in B’ **(D)** and in C’ **(E)** exhibit a peak value close to 1 at lag time 0. **F.** Comparison of CCF between pairs of PNs connected through neurites or somas shows no difference between these two situations (p>0.33, Mann-Whitney U, two-tailed, n=13 neurite pairs; n=14 soma pairs).

Furthermore, the correlated activity persists in presence of TCP (Figure 2A’), suggesting that it is independent of chemical synapses. While most of PN-PN pairs consist of cells in contact only through their neurites, others were in contact at the level of their cell bodies (Figure 2B-C). We evaluated whether the site of contact influenced the degree of correlated activity between PN-PN pairs in culture using a cross-correlation function (CCF). This approach describes the probability of coincidence detection of a calcium transient in one cell while a transient is observed in the second cell in the pair. CCF values close to one indicate a high coincidence of transients between two cells of the pair. The CCF analysis for PN pairs connected only by neurites (e.g., traces shown in Figure 2B’) show a strong correlation as maximal peak values were close to one at lag time 0 (Figure 2D). Similarly, a high correlation in spontaneous events is found in PNs connected through their soma (Figure 2C, 2C’, 2E, and F). No differences in the correlation coefficients between the two groups are observed (Figure 2F, P>0.33; Mann-Whitney U test), suggesting that the PN-PN coupled activity is independent of the site of contact between the cells. To determine if the correlated activity was specific to PN-PN pairs, spontaneous calcium transients in PN- GFP-negative pairs that were connected at the somas or through neurons in the same cultures were evaluated. There was little evidence of synchronized activity in between PNs and non-PN pairs, with correlation coefficients that are significantly lower compared to PN-PNs pairs (P<0.0001, Figure S1).

Taken together, these results demonstrate correlated spontaneous calcium activity between PN-PN pairs, that is independent of fast chemical neurotransmission and is dependent on PLTXII-sensitive voltage-gated calcium channels.

As the spontaneous coupled activity between PNs is independent of fast chemical synapses, we hypothesized that electrical synapses, mediated by gap junctions, might contribute to the correlated calcium activity. To test this idea, we directed a RNAi against *inx7* to AL PNs to decrease the expression of Inx7 specifically in this cell type. Knock- down of Inx7 expression is effective based on analysis of transcript in the knockdown flies (RNAi*^inx7^*; 85 ± 7% mRNA decrease as compared to UAS-RNAi*^inx7^* parental strain, n= 3 independent experiments). Cultures were prepared from brains of Inx7 knocked-down flies, and these contain PNs in physically connected pairs at a similar frequency as compared to cultures prepared from control flies (Figure 3A). In addition, the PNs found in these cultures generate spontaneous calcium transients like those detected in control cultures (Figure 3B). Nonetheless, in sharp contrast to control cultures, the calcium transients in PN-PN pairs expressing RNAi*^inx7^* show little or no correlated activity in representative calcium transient profiles (Figure 3B). Quantification of all PN-PN pairs expressing RNAi*^inx7^* evaluated demonstrates that decreasing Inx7 expression decreases correlated calcium activity in physically connected PN-PN pairs (Figure 3C). Overall, our data shows that Inx7 contributes to correlated calcium activity between cultured *Drosophila* AL PNs.

**Figure 3.**
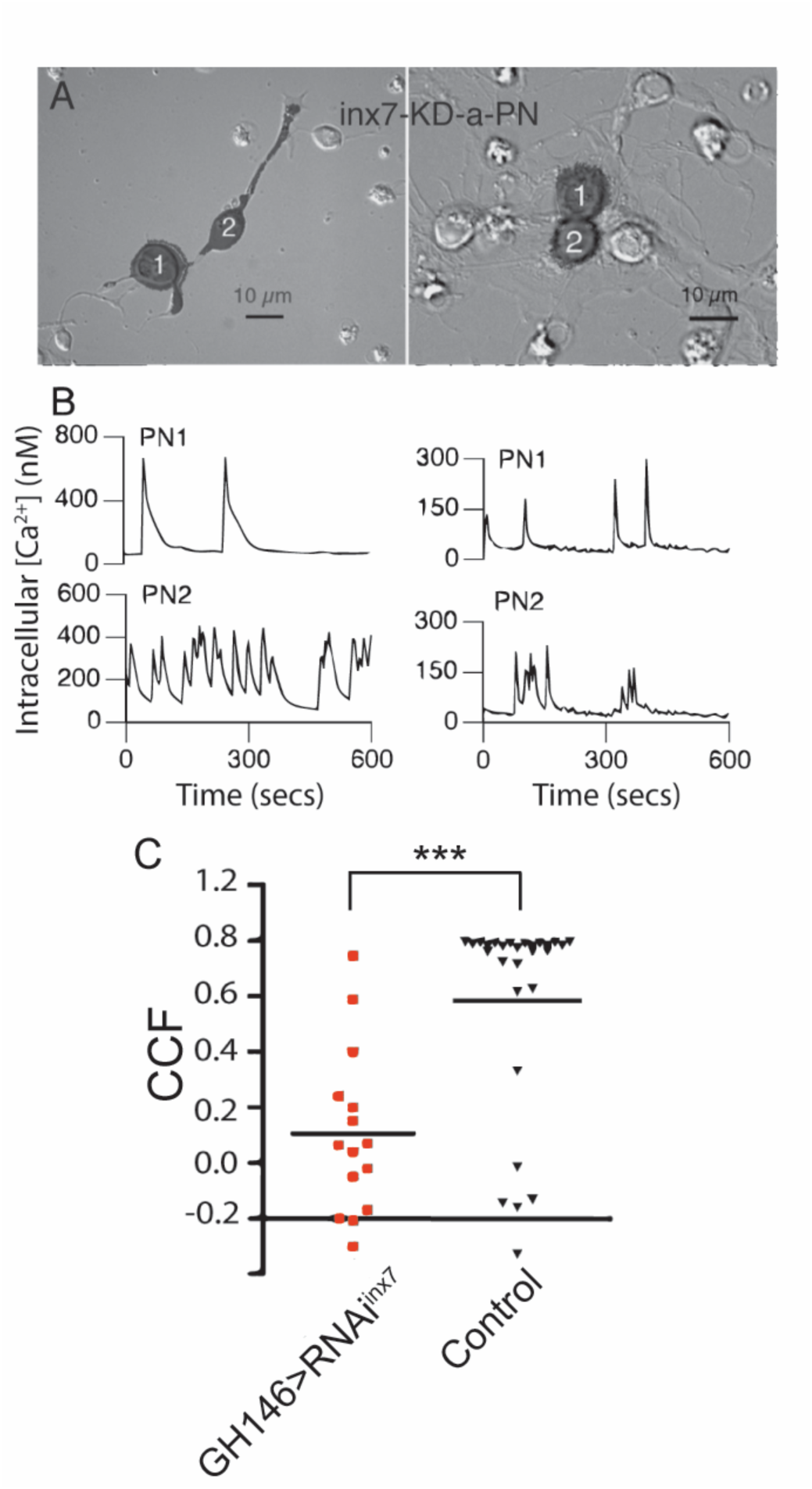
Knockdown of Inx7 in PNs reduces synchronous calcium activity observed in PN-PN pairs. **A.** Fluorescent mask (shaded black) projected on Nomarski image is shown for GFP positive PN-PN pairs expressing RNAi*^inx7^* in representative experiments. GFP-expressing cells are in contact through neurites (left) or cell bodies (right). **B.** Calcium transients recorded from PN-PN pairs expressing RNAi*^inx7^*in A. **C.** Correlation coefficients of all PN-PN pairs in neurons expressing RNAi*^inx7^* examined compared to all the PN-PN pairs in cultures made from control flies at the same time (***p<0.01; Kruskal-Wallis ANOVA, GH146>RNAi*^inx7^* n=15 pairs vs control n=28 pairs).

### Evoked calcium activity in AL PNs depends on Inx7

Correlated activity supports odorant processing in both the vertebrate OB and the insect AL [7, 20]. We evaluated whether Inx7 contributes to odor-evoked calcium responses in AL *in vivo*. Flies were exposed to different concentrations of vinegar (an attractive odorant) [11, 34] while neuronal activity was measured via calcium imaging in AL. Flies constitutively expressing the genetically-encoded calcium sensor GCaMP3 in the AL PNs (GH146,GCaMP3, Figure 4) were the control strain and were compared to flies recombinant for these transgenes crossed with flies bearing the UAS-RNAi*^inx7^* genetic element to knockdown the expression of Inx7.

**Figure 4.**
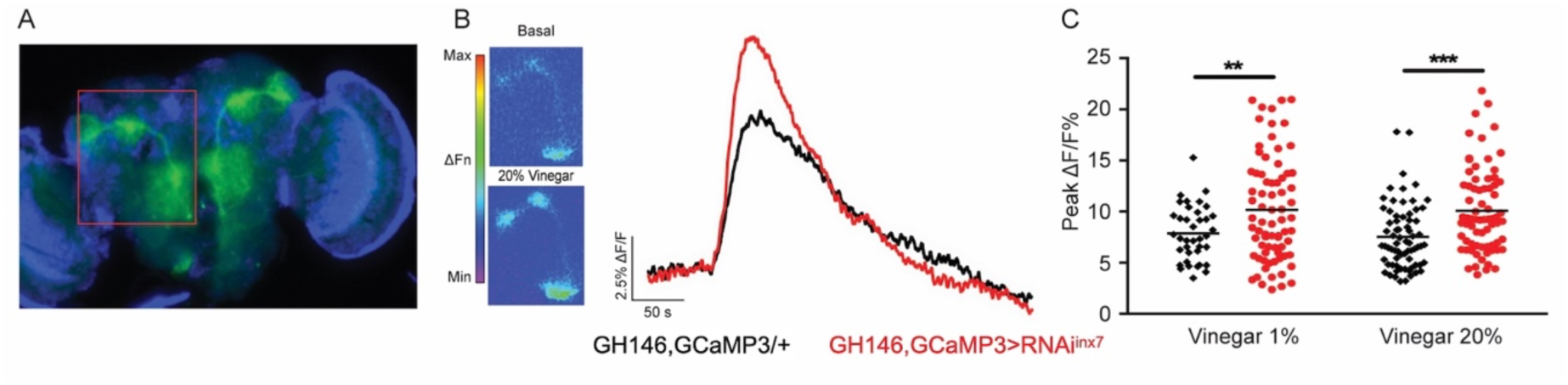
Inx7 knockdown affects vinegar-evoked calcium response in AL PNs. **A.** Fluorescent micrography showing the GCaMP3 expression confined to AL PNs using the Gal4 driver GH146. The red box shows GCaMP3 expression in AL glomeruli and projections to MB and lateral horn. **B.** Vinegar exposure increases intracellular calcium concentration in AL PNs seen as an increase in fluorescence. Representative traces of GCaMP3 fluorescence over time in AL glomeruli in response to 20% vinegar exposure are shown for control animals (GH146,GCamp3/+, black trace) and flies expressing RNAi*^inx7^* in AL PNs (GH146,GCaMP3>RNAi*^inx7^*, red trace). **C.** Peak calcium signals evoked by 1% and 20% vinegar in AL in control and RNAi*^inx7^*-expressing animals. An increase at both vinegar dilutions is observed upon Inx7 knock down. ** and *** indicate p<0.01 and p<0.001, two-way ANOVA followed by Sidak’s multiple comparisons test; n=38-74 events per genotype, recorded from 5 or more different flies per condition.

In response to the odor stimulus (20% vinegar), control flies exhibit a sharp increase in intracellular calcium in AL PNs, as previously described (Figure 4B, black trace) [34, 35]. Remarkably, in flies expressing the RNAi*^inx7^* in AL PNs, an augmented calcium response is detected compared to control animals (Figure 4B, red trace). The vinegar-induced increased calcium response is consistent across experiments using 1% and 20% vinegar dilutions (Figure 4C).

These results suggest that Inx7-dependent electrical synapses contribute functionally not only to synchronizing spontaneous calcium activity between PNs *in vitro* but also modulate neuronal activity in AL in response to odorant stimulus *in vivo*.

### Decreased Inx7 expression in PNs affects vinegar sensitivity and vinegar-evoked behaviors

The AL is an essential region responsible for the organization and processing of odor information before it is sent to higher centers in the *Drosophila* brain [3, 4]. To address whether the alteration in calcium activity in AL after Inx7 knock-down in PNs is important for the execution of behavioral responses to odorants, we used two behavioral assays.

First, we analyzed the naïve olfactory response to vinegar, an appetitive olfactory stimulus [34, 35]. Groups of starved flies were placed in a T-maze apparatus and exposed to 20% vinegar (in one arm) or fresh air (in the other). After a 2-min decision period, the number of flies preferring vinegar or air was quantified and a response index (RI) was calculated. This assay shows that down-regulation of Inx7 in PNs is sufficient to decrease by 50% the RI compared to control flies (Figure 5A).

**Figure 5.**
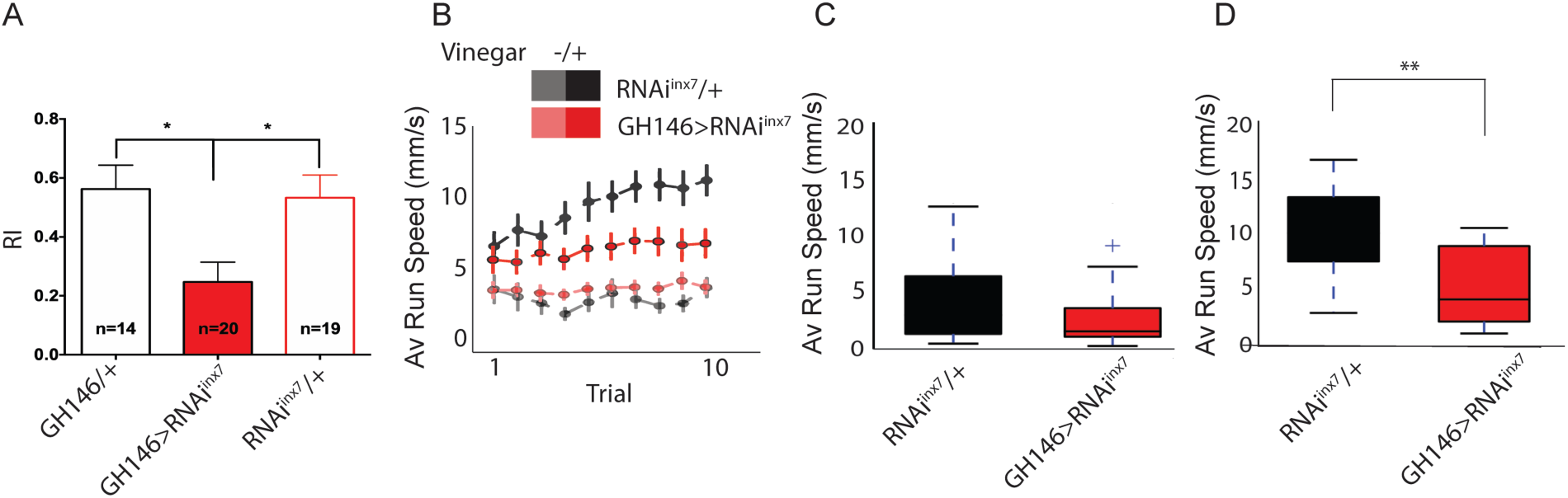
Behavioral responses to vinegar are affected in flies expressing a RNAi against *inx7* in AL PNs. **A.** Groups of 30-50 control flies (GH146/+ or RNAi*^inx7^*/+) show attraction towards vinegar, expressed as a positive index (RI) in a T-maze paradigm. Knocking down Inx7 in AL PNs (GH146>RNAi*^inx7^*) decreases RI. * indicates p<0.05; one-way ANOVA followed by Dunnett’s multiple comparisons test, n=number of flies tested, indicated for each genotype. **B.** Individual, starved, flies were acutely exposed to vinegar while the acute olfactive trail and locomotor behavior were recorded. This olfactory challenge was repeated consecutively ten times. The average running speed per trial is presented for control and RNAi*^inx7^*-expressing flies. An increase in this parameter is observed for the two strains, only when vinegar is presented in the control flies (black vs gray dots) and in the strain expressing RNAi*^inx7^*(red vs light red dots). As the number of trials increase, an escalation in the speed is observed in the control strain (black dots), an effect that is not observed in the RNAi*^inx7^*-expressing animals (red dots). No differences in the naïve locomotor response are observed in control or RNAi*^inx7^*-expressing flies as the number of trials increases (light red vs gray dots). **C and D** Comparison of the average speed for the 1^st^ trial (C) and 10^th^ trial (D). No difference between controls and RNAi expressing flies were observed in the first trial; however, differences were observed in the tenth trial **(D)**. ** indicate p<0.01, Mann-Whitney test.

We then evaluated the motivation of a fly for an appetitive stimulus [31]. To do this, we considered that flies display a preference for an appetitive cue, according to the integration of its internal state and the information provided by the cue. The integration between these two factors has been demonstrated in studies of appetitive memories, where it is reprted that flies generate memories only after a starvation period [36]. It has also been shown that starved flies persevere in seeking a food source when vinegar scent is repeatedly presented [31]. Thus, we hypothesized that alterations in calcium activity in RNAi*^inx7^* flies could result in impaired perseverance in seeking a food source. To assess this, we evaluated the locomotor behavior driven by appetitive food and behavior persistence in a spherical treadmill. Single starved flies were placed on top of a spherical treadmill and the average running speed towards the vinegar cue was quantified upon ten consecutive trials (Figure 5B). Under these conditions, both control and RNAi*^inx7^* expressing flies show strong attraction towards the first vinegar exposure, evidenced by an increase in speed of movement as compared to the situation when no stimulus was presented to animals. Importantly, no difference in the response is detected in the first trial in the two strains (Figure 5B-C). Interestingly, in this paradigm, the consecutive exposure to vinegar increases the speed movement in control flies (Figure 5B, RNAi*^inx7^*/+ group, black traces), a measure of increased motivation for food-seeking [31]. Remarkably, the flies expressing RNAi*^inx7^* in AL PNs show no increased motivation for food-seeking when vinegar is repetitively presented, as no change in speed of movement is observed across the consecutive trials (Figure 5B, GH146>RNAi*^inx7^* group, red traces). Differences arise between the two strains and were evident after the tenth trial (Figure 5D). The difference between the strains cannot explained by defects in basal locomotor performance in the RNAi*^inx7^* expressing flies, as naïve motor output is not affected in these flies when compared to control animals not exposed to the vinegar stimulus (Figure 5B, GH146>RNAi*^inx7^* and RNAi*^inx7^*/+ groups, light red and gray traces; Figure S2). This effect cannot be attributed to differences in the initial motivational state of RNAi*^inx7^*-expressing flies compared to controls either, since no differences between the two strains are detected in the first trial (Figure 5C). Similar results are obtained using an Inx7 null-mutant (Figure S3).

These data demonstrate that Inx7 plays a key functional role in AL PNs in processing olfactory information that guides appetitive behaviors in adult Drosophila.

## DISCUSSION

Synchronization of activity between cells plays an important role in the coordination of responses to different stimuli, in developing and mature olfactory systems. For instance, connexin36-containing gap junctions contribute to the synchronized activity between mitral and tufted cells in the OB, which is crucial for the organization, coordination and processing of sensory olfactory information in vertebrates [7, 8, 37]. Several reports show that gap junctions could play a similar role in the coordination of signals in the principal neurons of invertebrate AL, although the molecular determinants of such synchronization are not fully understood [20–22].

### Innexin 7 mediates PN-PN synchronized activity

*Drosophila melanogaster* has eight innexin genes, being ShakB (aka Inx8) the most extensively studied regarding its contribution to gap junction formation and electrical communication in CNS neurons [15, 16], particularly in the AL. For example, it is known that ShakB-containing gap junctions are necessary for correlated activity between PNs and excitatory LNs in the DA1 glomerulus in fly AL [21], and also for the synergistic response observed when flies are exposed to a pheromone and an odorant signaling food [11]. Moreover, it has been shown that homotypic PNs (i.e., neurons within the same AL glomerulus) can form electrical synapses in *Drosophila* AL, which are mediated by ShakB [21]. Whether other innexins expressed in the AL also play a role in the synchronized activity is not known.

Here, we have shown that Inx7 expression is required for synchronized calcium activity between physically connected PNs *in vitro*. This suggests that PNs possess the ability to establish electrical synapses with other PNs via gap junction proteins such as ShakB or Inx7. Supporting this notion, Phelan et al., 2008 [38] reported that ShakB expression is sufficient to promote gap junction formation between neighboring *Xenopus oocytes*. However, this would be mostly valid for homotypic PN pairs, since the number of PN-GFP negative pairs with high correlated calcium activity is smaller.

We detected a small number of PN-GFP-negative pairs that did show synchronized calcium activity. It is possible that the GFP-negative neurons in these pairs are unlabeled PNs since previous studies have shown that the GH146-Gal4 driver does not label all PNs [25] . Alternatively, previous reports suggest that LNs also interact with PNs through chemical, electrical, and mixed electro-chemical synapses, and that all these forms of neuron-to-neuron communication contribute to the olfactory network activity in the *Drosophila* AL [21, 22]. Thus, it is possible that in the culture system, the GFP-negative neurons correspond to AL LNs that in the fly brain are establishing gap junctions with PNs.

### Behavioral consequences of synchronized activity in AL

Electrical and/or calcium-synchronized activity have been associated with different functions in neurons and neural circuits [39–41]. Notably, calcium oscillations in the insect MB have been suggested to be relevant for olfactory learning and memory [42]. Further, spontaneous electrical activity has been shown in PNs in different invertebrates, where they seem to be modulating the processing of olfactory information [20, 43]. Although it is not clear the role of AL oscillations in odor processing, it has been proposed that they could contribute to guarantee the fidelity in the transmission of information within the olfactory circuit by refining the signal-to-noise ratio in downstream neurons, i.e., KCs or lateral horn neurons [20, 43].

To advance our understanding on the contributions of PN gap junctions to odorant processing in *Drosophila*, we carried out functional imaging experiments using vinegar as an appetitive odorant. Exposure to an appetitive stimulus resulted in an increase in calcium activity in PNs in control flies. Interestingly, the calcium response in RNAi*^inx7^*- expressing flies was higher when compared to control flies. Similar results have been previously reported: exposure to some odorants evokes an increase in the firing rate in PNs of the VC1 glomerulus, which is further increased in the *shakB^2^* loss of function mutant [21]. Thus, it is possible that expression of RNAi*^inx7^* in AL PNs causes a decoupling of PN-PN pairs that in turn increases calcium activity in these neurons. Our *in vitro* results does not support this idea, as calcium transients in AL PNs expressing RNAi*^lnx7^*are not different in frequency or amplitude from those recorded in single cultured AL PNs or in MB neurons *in vitro* [29]. Alternatively, it could be possible to propose that RNAi*^lnx7^* expression impairs recruitment of inhibitory information to PNs within different glomeruli. Impairment of inhibitory information into PNs could affect basal electrical properties of these neurons making them more excitable. This idea finds support in the description that inhibitory information is conveyed to PNs by LNs [44], and in the previous description of PN-LN electrical synapses being disrupted in mutants for the ShakB gap junction protein [20, 21].

The consequences of disrupting the Inx7-dependent gap junction in AL are evidenced as a reduced olfactory response towards the attractive stimulus vinegar. Furthermore, we demonstrated no change in the response to successive vinegar exposures in RNAi*^inx7^*-expressing flies. Functional integration of sensory information in an animal depends on different factors, such as the nature of the stimuli, the innate value of the information, and the inner state of the organisms, among others. Our data show that consecutive vinegar exposure results in behavior perseverance in control starved flies, as previously described [31]. The perseverance is observed as an increase in average speed in starved flies after each trial in which flies are exposed to vinegar. Such an increase in perseverance is not observed in RNAi*^inx7^*-expressing flies, while basal locomotor responses to vinegar in starved flies were indistinguishable between RNAi*^inx7^*- expressing and control flies. Thus, these data would suggest a role for Inx7-based electrical synapses at the AL PNs in olfactory detection, which could have further consequences in the integration of sensory information from the inner state of flies.

Overall, our results suggest that Inx7-based gap junctions are important contributors to the AL synchronized activity underlying olfactory processing and responses in adult Drosophila.

## Notes

### Competing Interest Statement

The authors have declared no competing interest.

