## supplemental Information for "The Innexin 7 gap junction protein contributes to synchronized activity in the *Drosophila* antennal lobe and regulates olfactory function"

**Supplementary Figures**

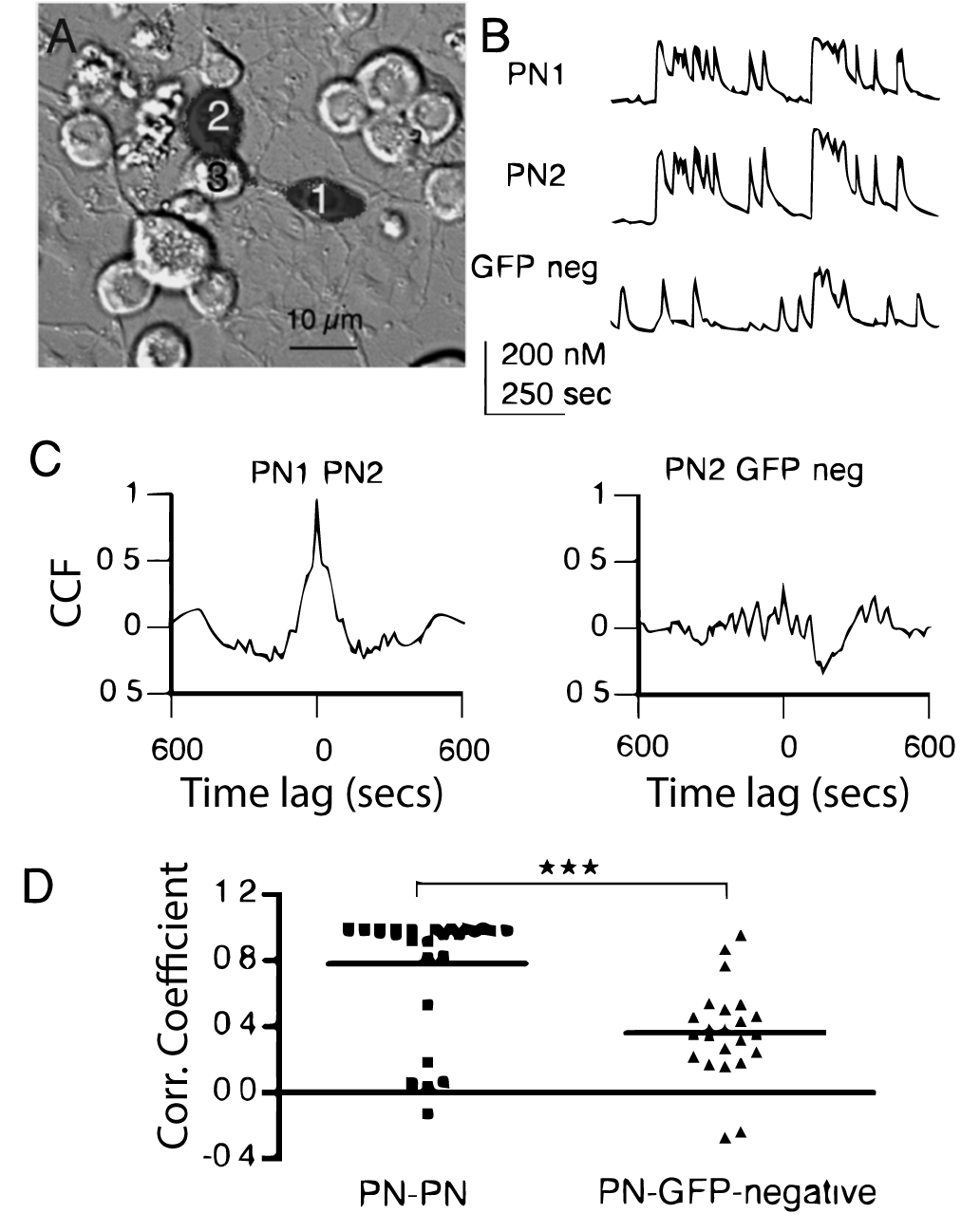

**Figure S1. PNs form connections that mediate correlated activity preferentially but not exclusively with other PNs. A.** PN pair (1 and 2) is in contact through neurites. A GFP-negative cell (3, GFPneg) is in contact with the cell body of PN2 and the neurites of PN1. Fluorescent mask (shaded black) projected on Nomarski image. **B.** Calcium transients in PNs show a highly correlated pattern of activity with each other but not with the GFP-negative neuron they are contacting. **C.** Cross correlation function analysis confirm a high degree of correlation between the PN1-PN2 pair with a value of 0.98 at lag time 0. In contrast the cross-correlation function for the PN2-GFP-negative pair has a value of .34 at lag time 0. **D.** Comparison of correlation coefficients for the PN-PN (n=28) and PN-GFP-negative (n=23) pairs illustrate the difference in mean (indicated by horizontal bars) and distribution. The PN-GFP-negative group has correlation coefficients that are significantly lower than the PN-PN group (***P<0.0001; Mann-Whitney U test, two-tailed). In 10% of the PN-GFP-negative pairs (2/23) the correlation coefficients are above 0.8 indicating that PNs can form connections that mediate correlated activity with other cell types.

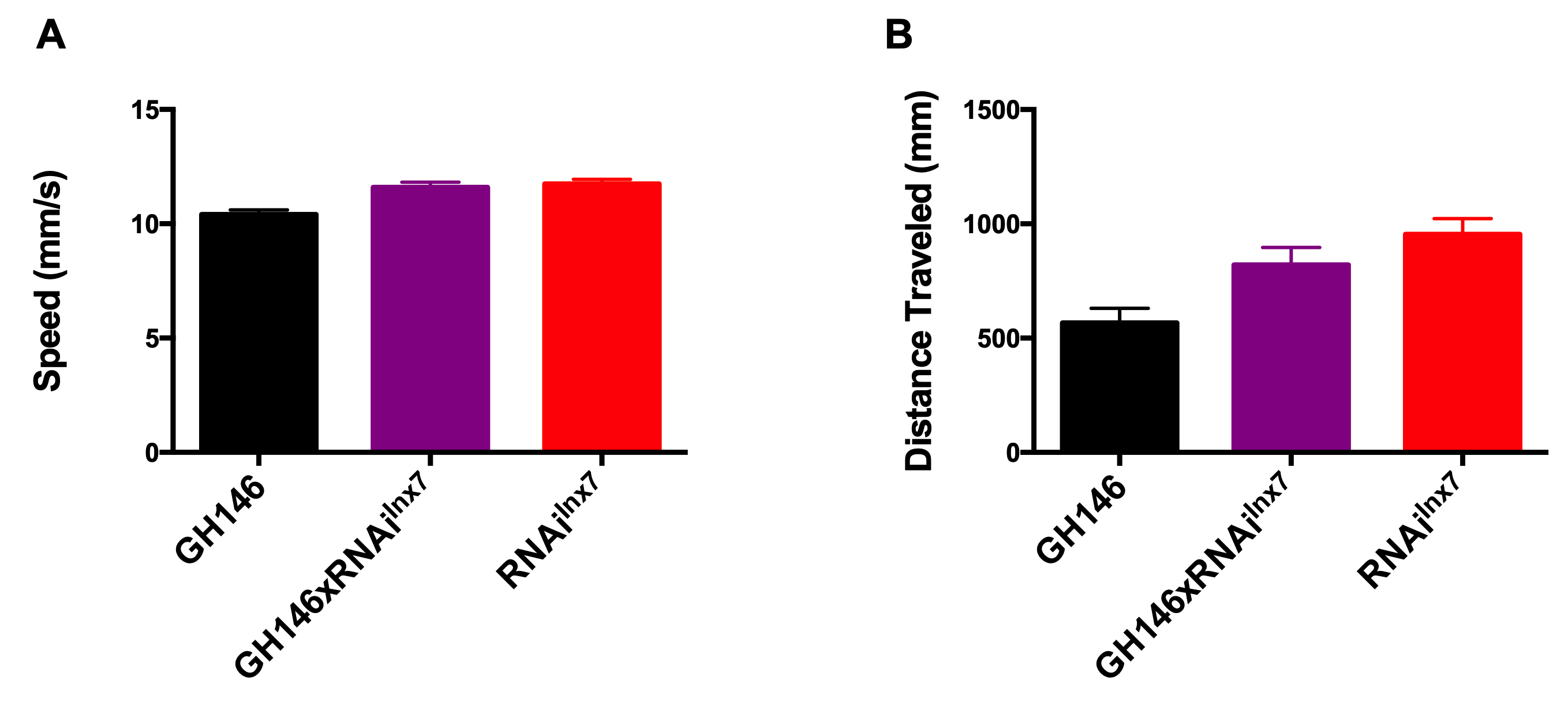

**Figure S2. Knockdown of inx7 in PNs does not modify the ability of flies to execute motor programs.** Different motor parameters were evaluated in animals expressing the RNAi^inx7^ in AL PNs. Here speed (A) and distance traveled (B) are shown. No statistical differences were detected in these parameters between RNAi^Inx7^-expressing flies and genetic controls. Data in A and B represents mean + SEM of at least 8 individual experiments. One-way ANOVA followed by indicates no differences between groups.

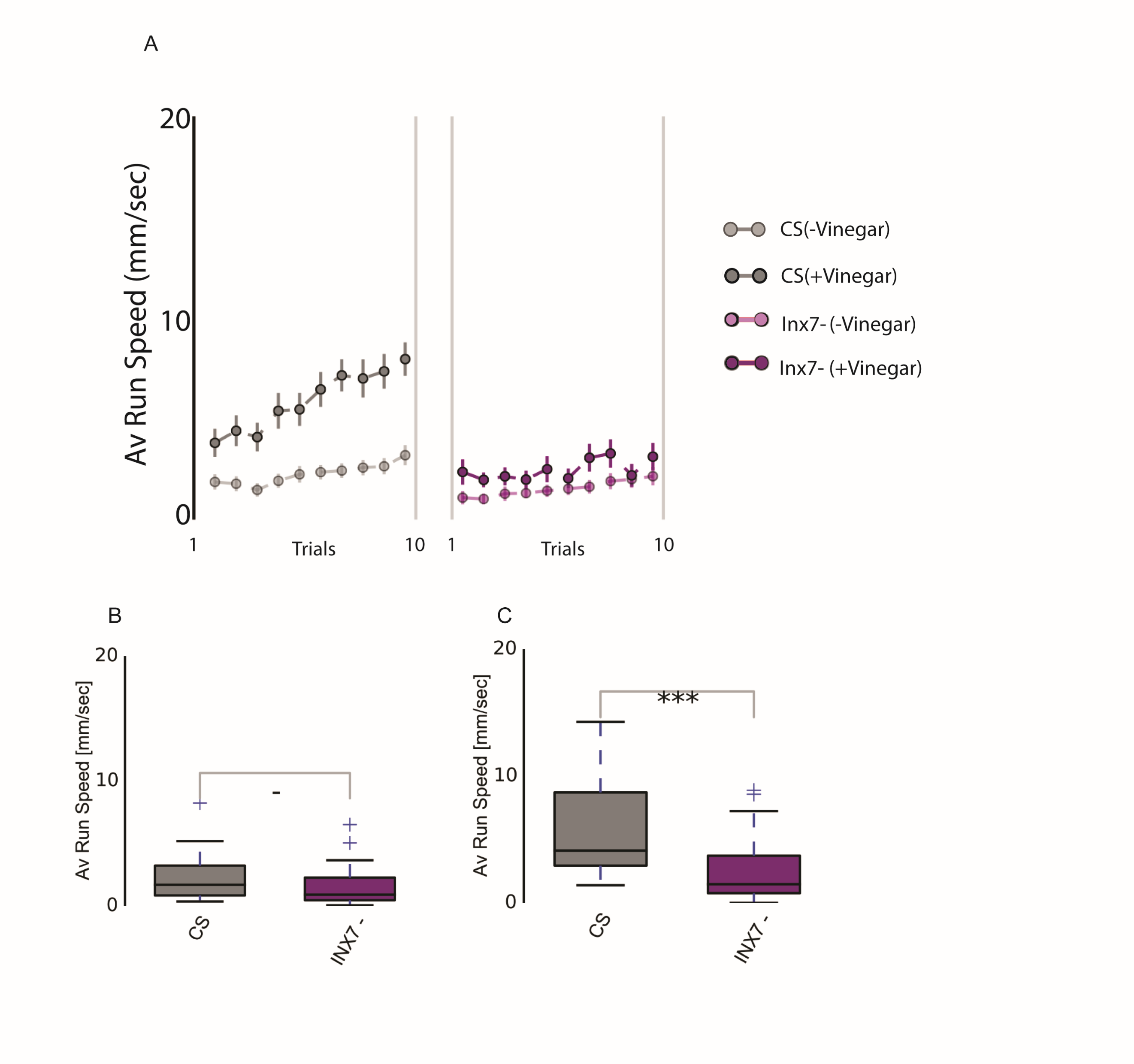

**Figure S3. Vinegar-evoked perseverance is affected in inx7 null mutants.** **A.** The average running speed per trial is observed for control (left panel) and inx7 mutant (right panel). **B.** Average running speed in first trial is not different in inx7 mutants as compared to control flies whereas the increase in motivation evoked by vinegar (i.e. increase in average running speed throughout the trials) is lost in the null mutants compared to controls (C). *** indicate p<0.01, Mann-Whitney test.
